# The Scalable Variant Call Representation: Enabling Genetic Analysis Beyond One Million Genomes

**DOI:** 10.1101/2024.01.09.574205

**Authors:** Timothy Poterba, Christopher Vittal, Daniel King, Daniel Goldstein, Jacqueline I. Goldstein, Patrick Schultz, Konrad J. Karczewski, Cotton Seed, Benjamin M. Neale

## Abstract

The Variant Call Format (VCF) is widely used in genome sequencing but scales poorly. For instance, we estimate a 150,000 genome VCF would occupy 900 TiB, making it both costly and complicated to produce and analyze. The issue stems from VCF’s requirement to densely represent both reference-genotypes and allele-indexed arrays. These requirements lead to unnecessary data duplication and, ultimately, very large files.

To address these challenges, we introduce the Scalable Variant Call Representation (SVCR). This representation reduces file sizes by ensuring they scale linearly with samples. SVCR achieves this by adopting reference blocks from the Genomic Variant Call Format (GVCF) and employing local allele indices. SVCR is also lossless and mergeable, allowing for N+1 and N+K incremental joint-calling.

We present two implementations of SVCR: SVCR-VCF, which encodes SVCR in VCF format, and VDS, which uses Hail’s native format. Our experiments confirm the linear scalability of SVCR-VCF and VDS, in contrast to the super-linear growth seen with standard VCF files. We also discuss the VDS Combiner, a scalable, open-source tool for producing a VDS from GVCFs and unique features of VDS which enable rapid data analysis. SVCR, and VDS in particular, ensure the scientific community can generate, analyze, and disseminate genetics datasets with millions of samples.

## Introduction

The pipeline for high-throughput sequencing involves a series of datatypes and the transformations between them:

1. *Sequencing* DNA to generate unaligned reads, often stored in a FASTQ.
2. *Aligning* to a reference genome to generate aligned reads, often stored in a BAM/CRAM.
3. *Variant calling* to generate genotype calls and metadata, often stored in a GVCF or VCF.
4. *Joint calling* to generate an analysis-ready genotype matrix, often stored in a PVCF.

*Sequencing*, *aligning*, and *variant calling* are straightforwardly sample-parallel, but *joint calling* is not. The latter necessarily combines multiple sample-oriented files into a single variant-oriented matrix. In this paper, we focus exclusively on *joint calling* and its data formats.

As cohort sizes grew so did the challenge of combining many sample-major GVCFs into one variant-major PVCF. These challenges have motivated the development of new tools (e.g DRAGEN gVCF Genotyper, GLNexus), new representations (e.g. msVCF, spvcf), new compressors (e.g. popvcf), and new formats (e.g. Savvy, Genomic Variant Store, GLNexus’s key-value store, GenomicsDB). These efforts draw from three approaches: sparsity of variant genotypes, sparsity of alleles, and amenability to sample parallelism. In this paper, we describe a framework which employs all three techniques to address the above challenges and yield exactly linear scaling of size in samples, sample parallelism in generation, incremental N+K mergeability, and a plaintext VCF representation.

### Formats and representations

The common representation of genomic sequences, Project VCF (PVCF), is untenable at scale. To see why, consider the addition of a single sample to a sequencing dataset with N samples. PVCF requires, for every novel variant in the new sample, an additional row containing one variant genotype and N homozygous reference genotypes.

Furthermore, each new allele that the N+1th sample introduces requires extension of existing fields to account for the new allele. For instance, the first N samples’ *allele depth* (AD) fields need a new “0” (no reads) entry for the new allele. Worse, the Phred-scaled genotype likelihood (PL) field grows quadratically in the number of alleles. The result is a repetitive and bloated representation, where the same uninformative value is stored over and over again, leading to inefficient use of storage and computational resources.

Throughout this paper we distinguish between a *representation* and a *format*. A representation, or data model, defines the expected set of fields, what they mean, and how they are related. A format describes a concrete implementation in terms of bytes. We similarly cleave the VCF Specification into the PVCF representation for cohort-level variant data and the VCF format for storing a variant-by-sample matrix of arbitrary data types in tab-delimited plaintext.

Specifically, PVCF defines the semantics of fields such as GT, AD, GP, PL, and, for list fields, the relationship between their length and the number of alternate alleles. VCF, as a format, describes, for example, how a number or a list is rendered in plaintext. The Variant Call Format supports representations other than PVCF: the single-sample Genomic Variant Call Format (GVCF) and the structural variant VCF (SV VCF). Analogously, the PVCF representation supports different formats: VCF and Binary Call Format (BCF).

### Variants scale with samples

When discussing the size of a dataset, we focus on the number of genotype records. The number of genotype records in a GVCF does not vary substantially across samples. We use *K* to refer to the average number of non-reference genotype records per sample, which, at 3-6 million, is much smaller than the number of bases in the human genome (3 billion).

In a multi-sample dataset, the expected number of loci at which at least one sample has a non-reference genotype is a function of the number of samples, *M(N)*. We call such loci *variant sites*, reserving the term *variant* to mean a particular allele at a particular locus. We use the term *genotype record* to refer to a genotype call and its associated quality metrics. By definition, *M(1) = K* and *M(N)*, for large *N,* approaches the number of loci in the reference genome. In contrast, the number of variants is unbounded due to the infinitude of possible insertion and deletion variants.

Consider *M(2)*. The second sample will share many variant sites with the first sample; however, both samples will have many unshared singletons. As a result, this second sample substantially increases the number of variant sites in the dataset. In contrast, the millionth sample will add significantly fewer variant sites because the previous 999,999 samples are likely to share many of its sites. Thus, *M(N)* is non-linear, which we confirm below.

PVCF represents a collection of sequences as a dense matrix, with one column per sequenced sample and one row for every variant site. PVCF permits both a *multiallelic* representation (wherein each locus appears in at most one row) and a *biallelic* representation (wherein loci are repeated such that each row contains one alternate allele). The number of genotype records in a multiallelic PVCF grows as

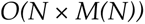

the number of genotype records in a biallelic PVCF grows as

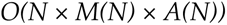

where *A(N)* is the expected number of alternate alleles at a variant site in a dataset with *N* samples.

Until variants are observed at all loci, each sample brings with it some number of new singleton sites. We empirically measured the relationship between the number of variant sites and the number of samples in subsets of the PVCF representation of the HGDP+1kG dataset. We found that the number of variant sites grows roughly with the square-root of N. Notice that we can measure the exponential relationship between M and N using a linear model in log-space.

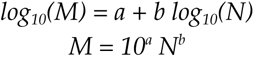

To explore this relationship, we generated multiallelic PVCFs from subsets of the HGDP+1kG dataset (Koenig et al. 2023). Figure 1a shows, for HGDP+1kG, *M ≈ N^0.46^*. Figure 1b shows the effect of this growth on the number of genotypes. The same experiment applied to subsets of gnomAD v2 found *M ≈ N^0.57^*. We stress that this is only evidence of an exponential relationship when *N* is between one and several thousand. Furthermore, this relationship depends on the ancestry mix of the dataset: diverse ancestries should increase the rate at which new variant sites are discovered. As the number of samples grows, the growth of variant sites must eventually decelerate, but not before sequencing datasets become impractically large.

**Figure 1.**
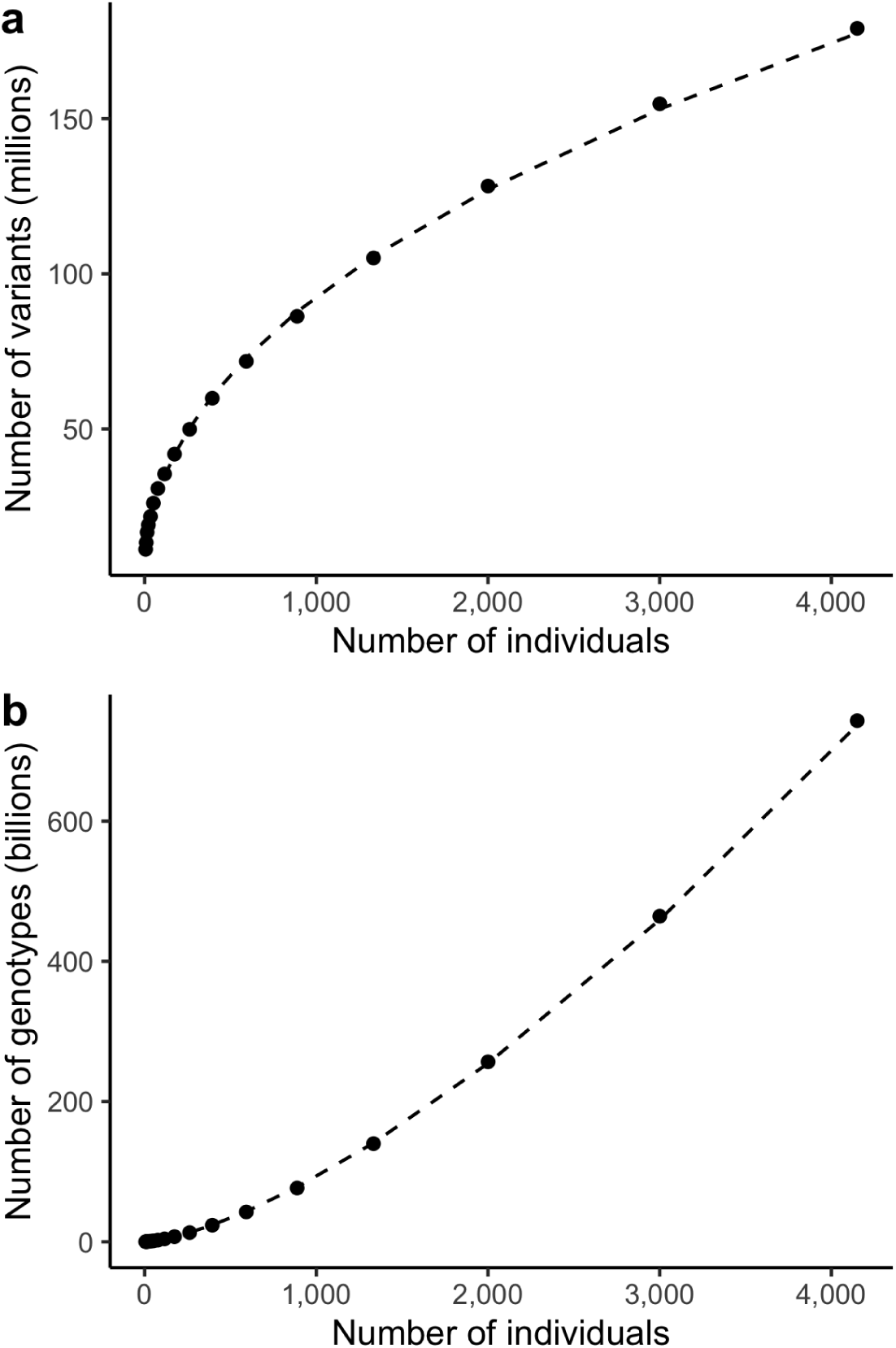
The number of variant sites (a) and total non-reference genotypes (b) scale with the number of samples. Each new sample brings new variant sites (a) which causes the dense matrix to grow super-linearly (b) in the number of samples. Dashed line indicates linear fit in log-log space, using all points with at least 100 individuals.

### Fields scale with alleles

In multiallelic PVCFs, not only does the number of genotype records grow super-linearly, but the size of each genotype record increases with the size of the cohort. In particular, sequencing datasets store per-allele quality metadata. For example, each genotype record stores an *allele depth (AD)* field containing one integer per allele observed at this locus.

This field grows linearly in the number of alleles at the locus even though most genotypes observe at most two alleles. Even worse, the *phred-scaled genotype likelihood (PL)* field stores the likelihood of all possible genotypes given the alleles at this locus. Within the autosomes, which are diploid, this field grows quadratically in the number of alleles. In practice, particularly in repetitive regions, a small fraction of loci have thousands to tens of thousands of alleles.

In summary, if *A(N)* is the average number of alleles per site, then the sum-total number of allele depths in a multiallelic PVCF file grows as,

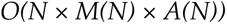

and the sum-total number of phred-scaled genotype likelihoods grows as,

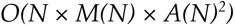

In practice, at a small number of sites, we observe thousands of alternate alleles. Sites like these are particularly challenging to analyze because of the quadratic growth of the PL field. We conjecture this quadratic growth explains why many PVCFs with more than a few thousand samples have dropped the PL field entirely or dropped it from multiallelic loci.

Notice that both multiallelic and biallelic datasets suffer from two forms of non-linear growth (genotypes and alleles). They differ only in where the growth appears. In the multiallelic representation, both genotype records and allele indexed fields grow super-linearly in the number of samples. In the biallelic representation, allele-indexed fields are bounded to length three but the number of genotype records grows even faster because highly multiallelic variant sites duplicate reference calls across every allele.

## Implementation

We propose a new representation for sequenced cohorts: the Scalable Variant Call Representation or SVCR. This representation is a generalization of GVCF to more than one sample. SVCR scales linearly in the number of samples. One or more GVCFs can be losslessly converted to a single SVCR dataset. Two SVCR datasets can be losslessly merged into a new SVCR dataset comprising samples from both. An SVCR dataset can be stored in many formats including VCF, Hail native format, and Google BigQuery.

### Overview

A dataset in the SVCR representation consists of zero or more variant sites. Variant sites are indexed and identified by genomic locus. There is at most one variant site per locus. Variant sites are multi-allelic. A dataset has a set of samples. Every variant site has a genotype record for every sample. A genotype record is either the unique missing value or a set of fields and their values. Any field value may be missing. Any value permitted for a VCF FORMAT field is permitted as a field value in SVCR.

### Column-sparsity

Within a single sample, SVCR stores adjacent reference genotype records as intervals, which are termed “reference blocks”. Each reference block has a locus interval, which must be contained within a single chromosome. A reference block is stored in the variant site identified by the interval’s first, or left-hand, locus. The size of the reference block, in base pairs, is stored in a genotype field named *LEN*. All reference block intervals for any particular sample must be disjoint.

Reference blocks are a form of column-sparsity by run-length encoding. At each locus, the genotype record of samples with a homozygous reference genotype is encoded in the underlying format as a missing value. The actual value is implicitly given by that sample’s overlapping reference block. Preservation of the reference blocks enables the combination of two or more SVCR datasets into a new SVCR dataset.

### Local alleles

Consider a highly polymorphic low-complexity insertion-deletion locus with 1,000 alternate alleles in a PVCF with 100,000 whole genomes. The VCF spec mandates that the allelic depth field *AD* has one value per allele including the reference (*R-numbered*), so AD records at this site must contain 1,000 elements denoting the depth of reads supporting each allele. For any particular sample, it is likely that at most one or two of these alternate alleles are observed. In the PVCF representation, the *AD* field for such a sample would contain between 998 and 999 zeros. The GVCF representation, in contrast, contains at most three entries. As described previously, this problem is significantly worse for fields like Phred-scaled genotype likelihoods (*PL*), which have one value for each possible genotype configuration (*G-numbered*). Data generators have mitigated this problem with approaches such as truncating the list of alternate alleles at any given locus to a maximum number, or disseminating PVCFs with only genotype (*GT*) fields and no quality metadata. Each of these approaches has drawbacks: truncating alternate alleles removes potentially pathogenic information from the dataset and masks information from short tandem repeats (STRs), and dropping all quality fields makes quality control much more difficult.

SVCR introduces a new entry-level field, *local alleles (LA).* At each locus, for each sample, an LA field indicates which alleles were observed in this sample. The *global alleles* refers to the set of alleles observed in any sample at this locus. Concretely, the LA field is an injective function from local allele indices to global allele indices (Figure 2). Since the number of local allele indices is finite (in a diploid genotype, there are two), we represent the LA field as an array. For example, at a locus with 1,000 global alleles, an LA value of [0, 4, 5] indicates that the first local allele corresponds to the zeroth (or reference) global allele, the second local allele corresponds to the fourth global allele, and the third local allele corresponds to the fifth global allele. For convenience, we require the first element of LA to always be zero.

**Figure 2.**
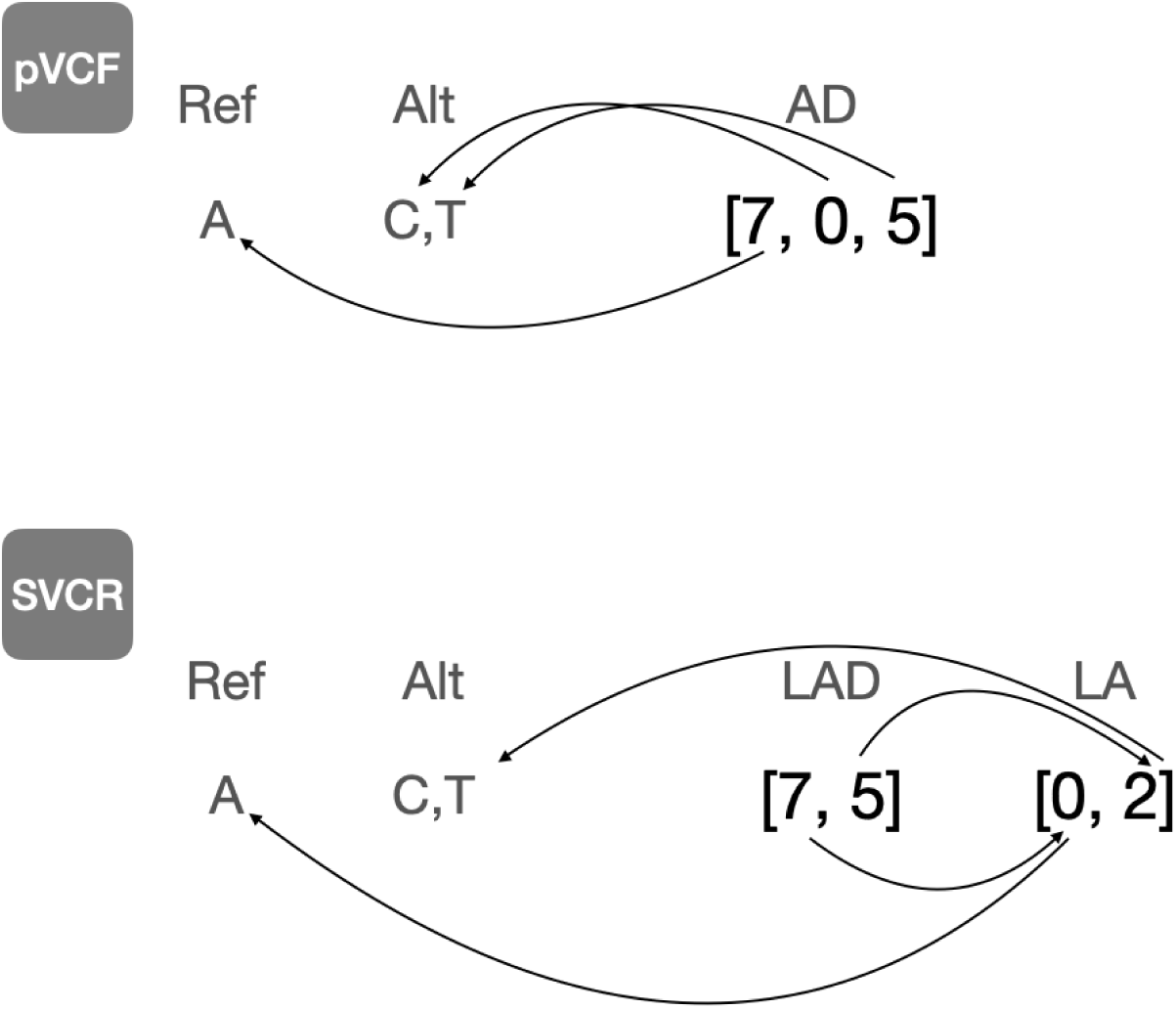
Local alleles. The AD field is an array field containing one element per allele, including the reference. The AD field entries indicate, for each allele, the number of informative reads. The arrows terminating at an allele indicate to which allele each value corresponds. The LAD field is a locally indexed array field storing the same data. Note that we need not store information for the unobserved C allele, and that the two-step paths lead to the same alleles as the one-step path.

For each genotype-level allele-indexed field from the VCF spec, we define a new field whose name begins with “L”. The field only contains the elements corresponding to this sample’s local alleles. Consider a variant site with five alleles, A, AA, AAA, AAAA, AAAAA, and a sample with the genotype AA/AAAA with 14 reads supporting AA and 16 reads supporting AAAA. The local alleles array will be [0, 1, 3], the allelic depth will be [0, 14, 0, 16, 0], and the local allelic depth array will be [0, 14, 16].

Although the GT field is not problematic, we strongly recommend using an LGT field in SVCR. In particular, when removing a sample, an implementation of SVCR may wish to remove global alleles which no longer appear in any samples. Removing a global allele requires modifying every local alleles entry *and also* updating any globally-indexed fields, such as the GT.

### SVCR-VCF

We encode SVCR in VCF by requiring two FORMAT fields: *LA*, an array of integers, and *LEN*, an integer (described above). Reference blocks must have a non-missing *LEN*, should have a non-missing *DP* & *GQ*, may have a non-missing *LGT*, and should have a missing *LA*, *LPL*, and *LAD*. When a sample has an overlapping reference block at a locus, all of that sample’s FORMAT field values must be missing. Non-reference genotypes must have a missing *LEN*, must have a non-missing *LA*, and should have a non-missing *DP*, *GQ*, *LGT*, *LAD*, and *LPL*.

SVCR introduces three new settings for the “Number” of a VCF list field:

- *LOCAL-A* numbered fields should have one list element for each allele present in *LA* minus one (to exclude the reference). For example: *LEC*.
- *LOCAL-R* numbered fields should have one list element for each allele present in *LA* (including the reference). For example: *LAD*.
- *LOCAL-G* numbered fields should have one list element for each possible genotype depending on the number of alleles N_LA_ and the ploidy of the genotype call. If diploid, (N_LA_ * (N_LA_+1)) / 2. If haploid, N_LA_. For example: *LPL*.

Figure 3 demonstrates the equivalent representation of twenty base pairs of two samples in GVCF, PVCF, and SVCR-VCF.

**Figure 3.**
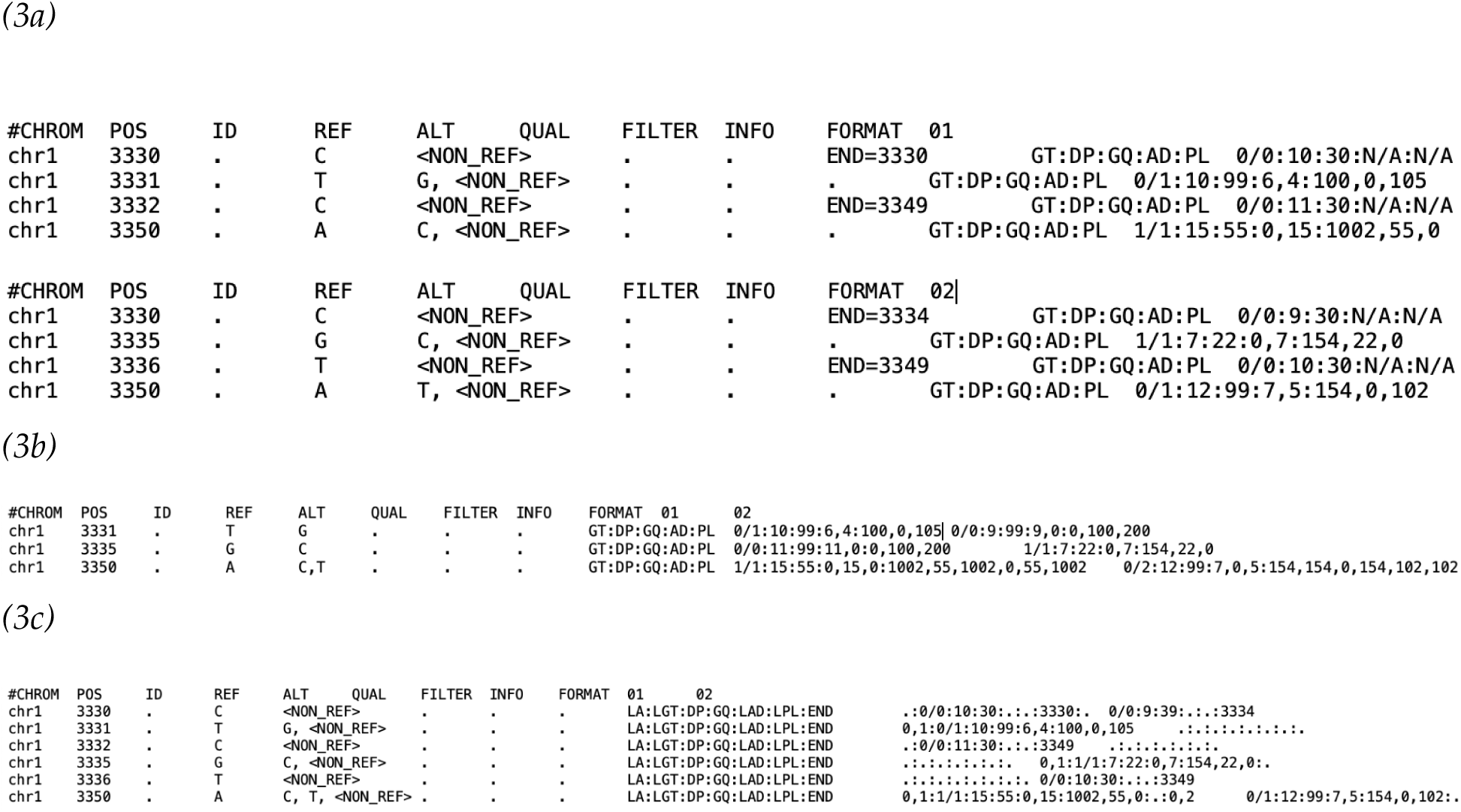
Representing sequences. In each panel, two different samples, 01 and 02, are shown in different representations, including a) two GVCF files, b) a PVCF file, and c) an SVCR-VCF file.

**Figure 4.**
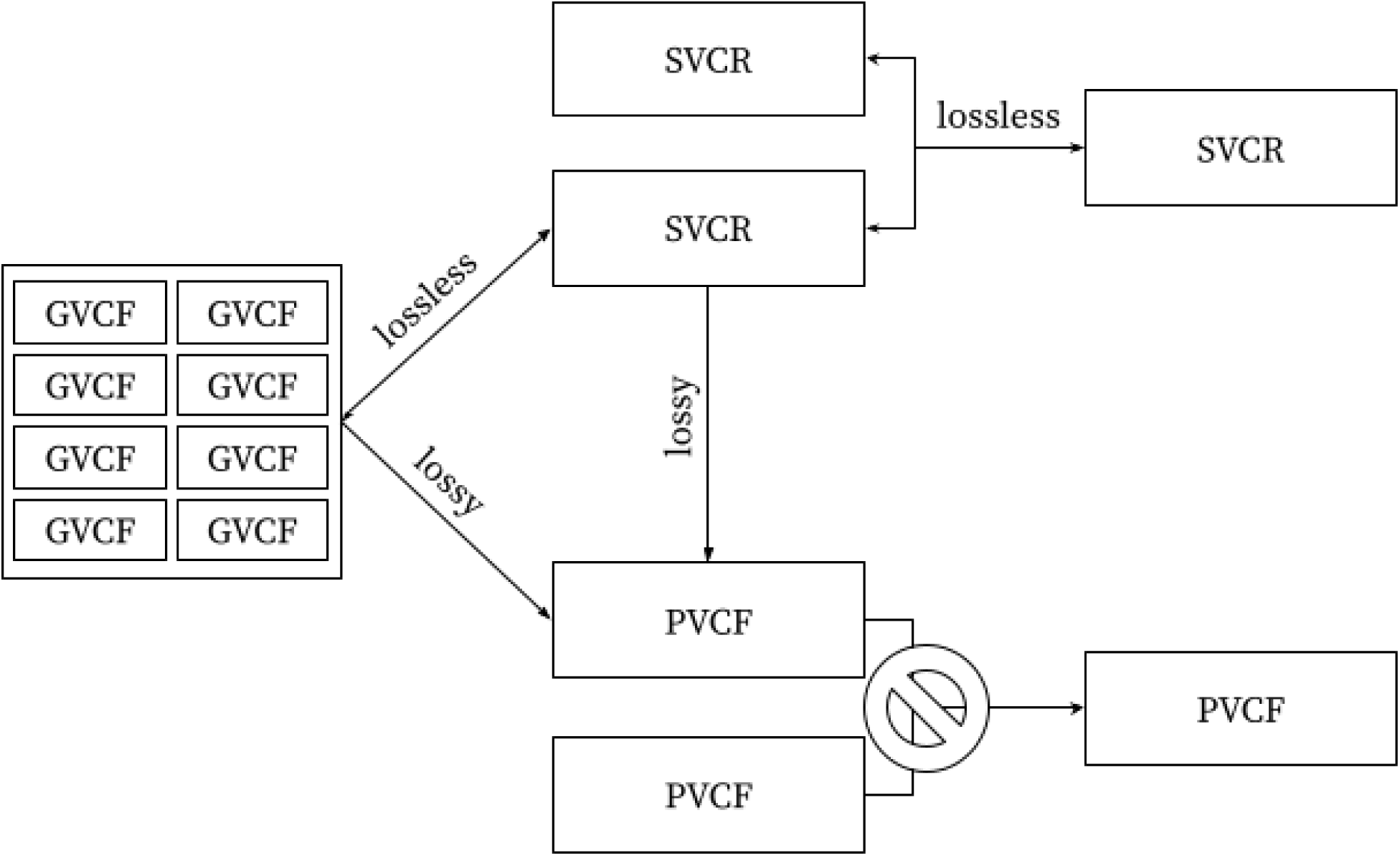
The graph of interconversion and combination operations. Notice that PVCF is a sink: crucial information is lost when producing a PVCF—missing data & homozygous reference calls can not be unambiguously resolved if the quality metrics at non-variant sites are discarded.

### VDS

We also encode SVCR in a Hail native format called Variant Dataset or VDS (unrelated to the Hail 0.1 VariantDataset except in name). A VDS comprises two Hail matrix tables. One Hail matrix table contains reference data the other contains variant data. The reference matrix table has rows keyed by locus, columns keyed by sample identifier, and exactly three entry fields: *LEN* (*int32*), *DP* (*int32*), and *GQ* (*int32*). Entries which are not the start of a reference block are missing values.

The variant matrix table has rows keyed by locus and alleles, columns keyed by sample id, and at least two entry fields, *LA* (*array<int32>*) and *LGT* (*call*), and usually also has *LAD* (*array<int32>*), *LPL* (*array<int32>*), *DP* (*int32*), *GQ* (*int32*), and *gvcf_info* (*struct*<…>). The latter contains all the GVCF INFO fields for that sample.

For data interchange, we recommend a format that uses a single file. For analysis, we recommend storing reference and variant data in two files for three reasons:

1. The schemas for reference and variant records differ; therefore, a combined representation pays the overhead of mixing these schemas. We observed a roughly 10% size reduction in both VCF and VDS from a split format.
2. The split representation makes it easier to filter variants without inadvertently removing reference blocks.
3. Analytical and quality control methods that operate on the smaller variant-only table are faster and cheaper because they read substantially less data. For example, the newly-developed CHARR (Lu et al., 2023) achieves substantial speedups as it only needs to access variant data (homozygous genotypes).

### Sparsity in representation versus sparsity in format

It should be noted that while SVCR is sparse—only a small number of matrix entries contain information—both the GZIP-compressed VCF and Hail VDS realizations of SVCR contain high numbers of explicit “missing” sentinel values. Both formats use compression techniques which rapidly & efficiently compress and decompress these repeated values, reducing the gain from switching to an explicitly sparse format. We anticipate substantial reductions in analysis cost and time from a system that stores *and computes* directly on the sparse representation.

### Densification

The reference information for all samples at some locus *L* is not contained within a single row of the matrix at *L*. Instead, this information is stored in overlapping reference blocks that appear at their start location, which is necessarily some previous locus. We call producing a single row containing all reference and variant information for all samples at a particular locus *densification*. In general, densifying a locus requires searching all previous loci until an overlapping reference block for every sample has been identified.

### Mitigation by periodic reference checkpoints

We propose two strategies to bound the size of each reference block and thus bound the search distance. Reference blocks could either be split at some fixed size limit or at fixed intervals over the genome. Both strategies add reference blocks and therefore trade increased space for reduced densification time. Both strategies have a parameter *K* such that, at a particular variant site, for all samples, all overlapping reference blocks begin no further than *K* base pairs prior.

#### Splitting at a fixed period

Split reference blocks at every *K* bases. Store reference blocks for all non-variant samples at each split location. If this dataset is also partitioned into *K*-sized partitions, the information necessary to realize any partition’s dense sub-matrix of genotypes is fully contained in that partition. This approach inflates the reference data by splitting every reference block that spans the *K-*th locus even if that reference block is already tiny. As shown below, this inflation relative to the next strategy is very modest.

#### Splitting at a size threshold

Split reference blocks spanning more than *K* bases into two or more reference blocks spanning at most *K* bases. For instance, with a value of *K=10,000* a reference block starting at *chr1:4500* with length *38000* would be split into reference blocks

#### chr1:4500-14499, chr1:14500-24499, chr1:24500-34499, chr1:34500-42499

This strategy introduces the minimum number of new reference blocks to ensure the *K* invariant; however, in partitioned datasets, data from previous partitions may be necessary to realize the dense sub-matrix of genotypes at any partition after the first.

#### K is a parameter of the dataset

We do not mandate or recommend any particular value of *K* because the length distribution of reference blocks substantially depends on experimental design, variant calling, and reference block bin resolution. For example, whole exome sequencing produces shorter reference blocks due to the fluctuations in depth from partial capture of the genome. In contrast, high coverage whole genome sequencing can produce long blocks of similar-quality reference calls in regions of constant depth. Table 1 and Table 2 present the measured number of reference blocks after splitting the HGDP+1kG dataset using each strategy.

**Table 1.** Data size is inversely proportional to the period when splitting at a fixed period. Using this strategy, a period of 5,000 bases introduces a modest 0.15% increase in size. The increased sizes of other periods are also shown. At a period of 50,000 bases, only a few hundred reference blocks are added.

| *K (fixed period)* | Billions of Reference Blocks after Splitting | Ratio of Split to Unsplit Reference Blocks |
| --- | --- | --- |
| 50,000 | 73.1 | 1.0000 |
| 10,000 | 73.1 | 1.0002 |
| 5,000 | 73.2 | 1.0015 |
| 1,000 | 77.7 | 1.0634 |
| 500 | 87.1 | 1.1925 |
| 100 | 175.5 | 2.4013 |

**Table 2.** Data size is inversely proportional to the threshold when splitting at a size threshold. In practice, compared to splitting at a fixed period, this strategy is only mildly more parsimonious.

| *K (maximum size)* | Billions of Reference Blocks after Splitting | Ratio of Split to Unsplit Reference Blocks |
| --- | --- | --- |
| 50,000 | 73.1 | 1.0000 |
| 10,000 | 73.1 | 1.0002 |
| 5,000 | 73.2 | 1.0015 |
| 1,000 | 77.7 | 1.0634 |
| 500 | 87.1 | 1.1922 |
| 100 | 175.3 | 2.3985 |

In our experience, the granularity of reference block GQ bins has a large impact on the average length of reference blocks in a given dataset. Whole genome sequencing data also tends to have longer reference blocks than whole exome data, due to the absence of fixed capture boundaries and more consistent coverage across the genome.

### Streaming densification

The previous discussion and mitigations address reading one particular locus. We call this a “point-query”. In contrast, applications that read all or most of the dataset do not need these mitigations. Instead, streaming densification keeps one overlapping reference block per sample, swapping in new reference blocks as they appear in the stream of records. As stated above, this dense matrix is super-linear in size. Although a streaming application does not reify the entire matrix in memory, if it performs computation on each dense entry, the compute-time, and thus cost, will grow super-linearly. For example, computing the allele frequency necessarily inspects averages over every call, even the homozygous reference calls. A clever implementation would avoid densification entirely.

### GVCF is a single-sample SVCR

SVCR is a multi-sample generalization of GVCF. Indeed, an SVCR with one sample is equivalent to a GVCF. In a GVCF, the global alleles and the local alleles are the same because there is only one sample. In this setting locally-indexed and globally-indexed fields are equivalent. Specifically, in a GVCF, the *AD* and *PL* fields must be equal to the *LAD* and *LPL* fields. The lossless transformation between a GVCF and SVCR-VCF is as simple as (1) adding a field LA to create an identity function of the alleles, (2) adding an “L” prefix to the name of allele-indexed fields, (3) adding a *LEN* field to the FORMAT derived from the position and the *END* field.

### Hierarchical joint calling

An SVCR may be produced in a tree-structured manner because it is combinable with itself and with GVCFs. This allows some sample-wise parallelism in the production of an SVCR dataset from GVCFs. Suppose we have 100,000 GVCFs. We may, in parallel, combine 1,000 groups of 100 GVCFs each into 1,000 SVCR datasets. We may then, in parallel, combine ten groups of 100 SVCR datasets into ten SVCR datasets. Finally, we may combine the ten SVCR datasets into one SVCR dataset. This entire process is also fully variant parallel.

## Results

### Joint calling & analysis of nearly one million exomes

SVCR has been used since 2019 to merge, store, and analyze large sequencing cohorts. The Hail library includes a “VDS Combiner” that can combine zero or more VDSes with zero or more GVCFs into a single VDS in a parallel and fault-tolerant manner. Five large datasets produced by the VDS Combiner directly from GVCF files are:

1. gnomAD v3.1. 153,000 genomes. 2019. (Chen, et al. 2020)
2. Center for Common Disease Genomics (CCDG) exomes. 203,000 exomes. 2021. (Felsenfeld 2018)
3. CCDG genomes. 136,000 genomes. 2021. (Felsenfeld 2018)
4. gnomAD v4. 955,000 exomes. 2021 (Chao, et al. 2023).
5. Blended Genome Exome (BGE) Wave 1. 82,000 exomes. 2023. (Howrigan 2022)

The All of Us research project also used the VDS Combiner, but started from tens of Avro files each containing 4,000 complete samples in a GVCF-like representation.

6. All of Us, April 2023 data freeze. 245,350 genomes. (All of Us, 2022)

In addition to producing the VDS for each of these datasets, the Hail system was used to perform quality control and analyze each dataset.

The VDS Combiner is implemented in Hail Query, an open-source, partitioned, horizontally-scalable, spot-tolerant, dataframe system and genomic analysis library with a Python API. Hail Query, and therefore the VDS Combiner, can run on any Apache Spark or Hail Batch cluster. A managed version of the former can be found in all the major clouds including AWS, Google, Azure, and Alibaba.

Hail Query is pervasively designed to support spot instances. At the time of writing, the cost of a spot core as a fraction of the non-spot price of general purpose families in AWS was about 40-50%; in GCP, about 21-40%; and in Azure, about 10-25%.

Unlike other joint calling systems, Hail Query is also a general purpose analysis system supporting relational algebra (e.g. filter, aggregate, group-by, order-by), distributed linear algebra (e.g. PCA, matrix multiplication), and export to many formats (e.g. BGEN, PLINK, VCF, TSV). The Hail Query language, which is used to manipulate values within datasets, supports a wide variety of data types (e.g. integers, floating-point numbers, strings, genomic loci, genotype calls, arrays, sets, dictionaries, tuples, ndarrays, and structs) and a wide variety of operations on these types (e.g. random functions, statistical distributions, statistical tests, linear and logistic regression, linear algebra, collection iteration & aggregation).

### Size in bytes

Consider subsets of the HGDP+1kG dataset in PVCF, SVCR-VCF, and VDS. The total size grows super-linearly with sample size for PVCF but linearly for SVCR-VCF and VDS (Figure 5a). Further confirming this, we see the size per sample increases for PVCF, while approaching an asymptote for SVCR-VCF and VDS (Figure 5b), as expected since each human genome contains an approximately fixed amount of information.

**Figure 5.**
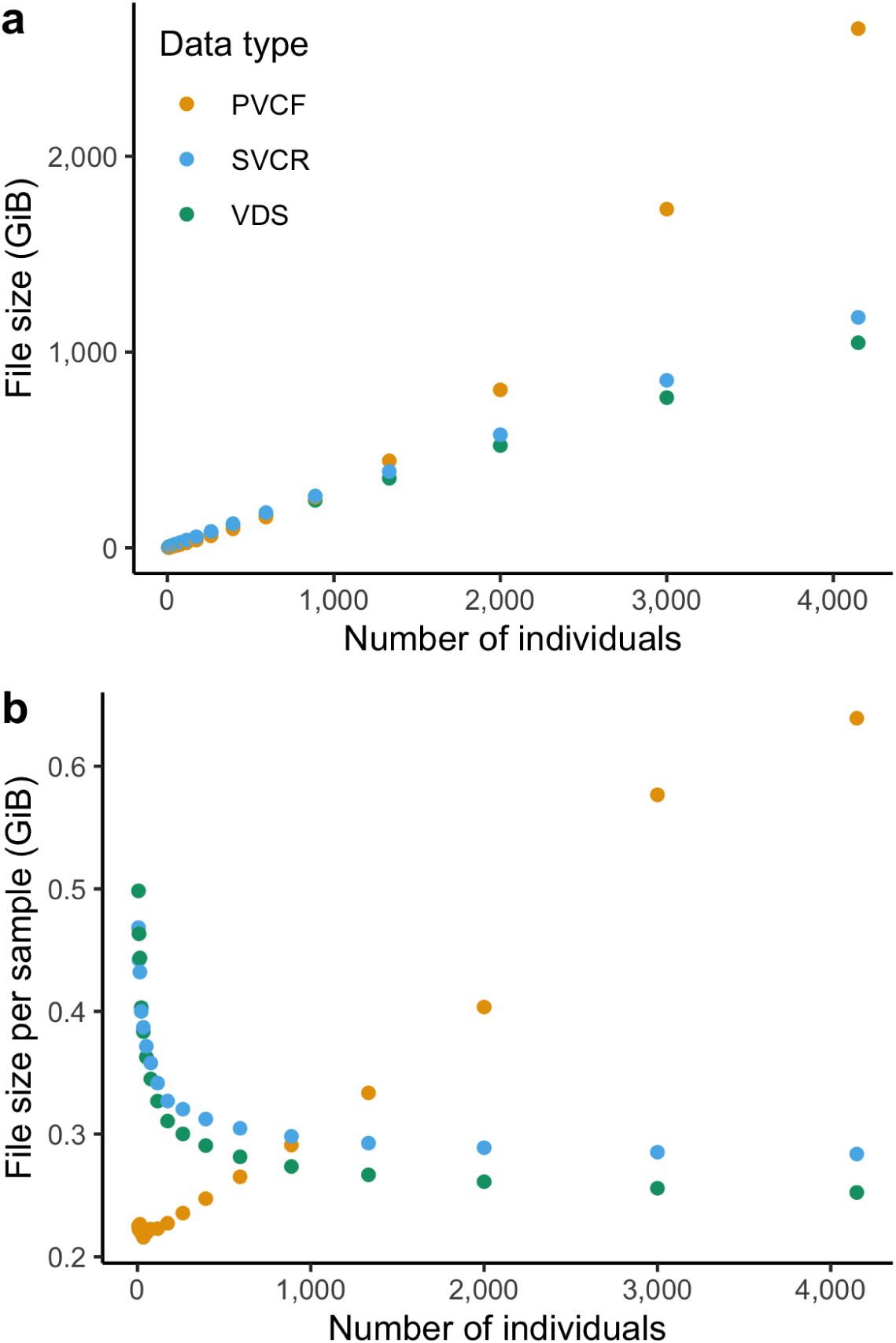
Scaling of dataset size of sequenced whole genomes represented in Hail LZ4-compressed VDS, gzip-compressed SVCR-VCF and gzip-compressed PVCF. In (a) total size in Gibibytes is plotted on the y axis, and in (b), plotting the size per sample (total size divided by number of samples) provides a clear view of the super-linear scaling of PVCF.

It is critical to note that exact figures of size per sample should not be used to compare representations and formats without additional context that reflects the granularity and type of information contained. For example, choices in resolution of reference blocks and the selection of included FORMAT fields can lead to a substantial size difference in single-sample GVCFs for the same sequencing experiments, and these ratios are preserved in SVCR.

We did not systematically generate PVCFs or SVCR-VCFs for larger sample sizes due to the high cost of these experiments, but we show that this asymptotic trend persists in a VDS at larger cohort sizes with subsets of gnomAD (Figure 6), and list a few selected examples of PVCF and VDS size in Table 3.

**Figure 6.**
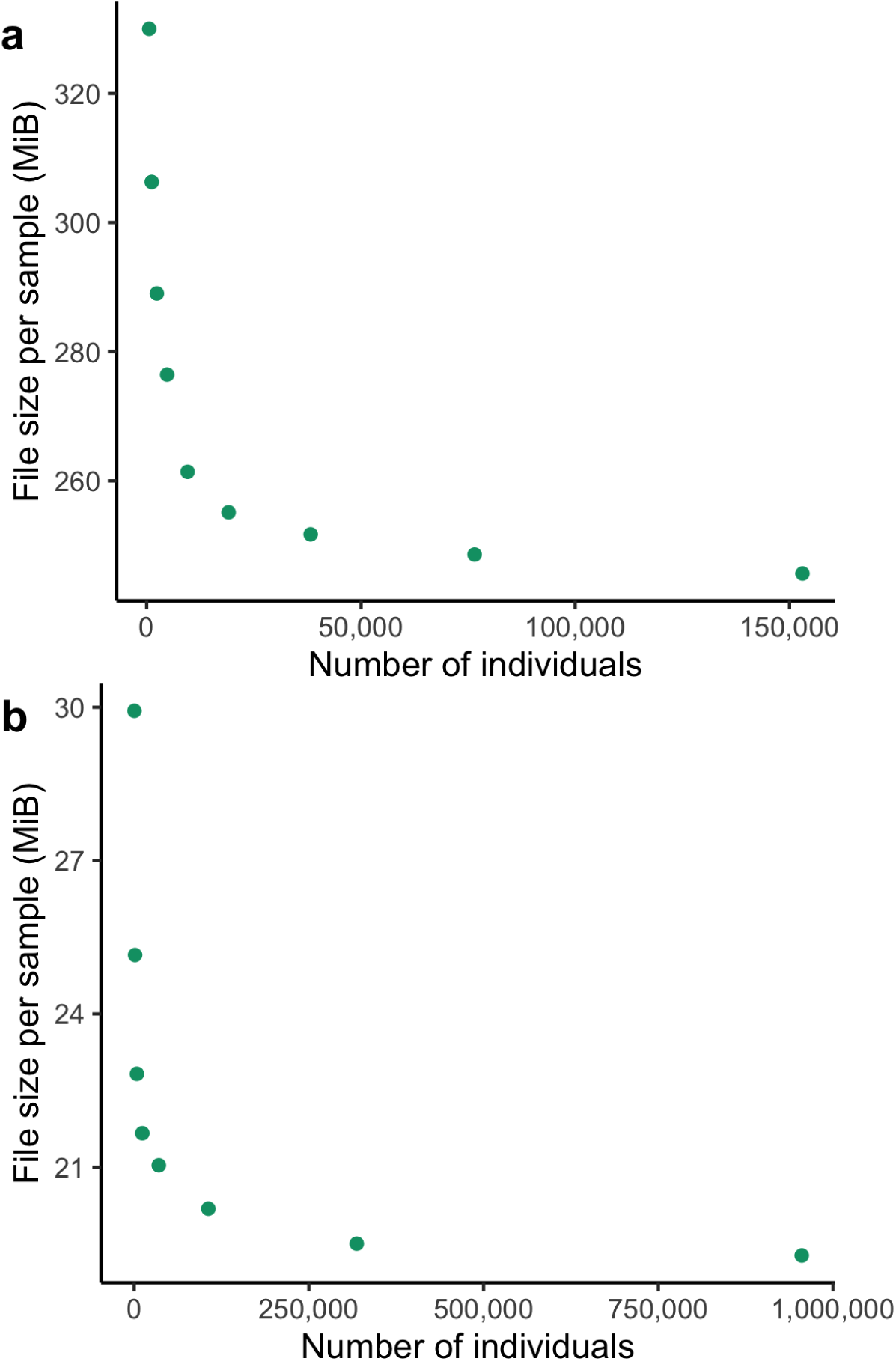
The growth of size in bytes <u>per sample</u> of subsets (a) gnomAD v3.1 exomes and (b) of gnomAD v4 exomes represented as VDS.

**Table 3.** Sizes and size per sample of selected large callsets. A blank PVCF cell indicates that a PVCF was never generated. A blank GVCF cell indicates the sum-total size of the GVCFs is no longer available. The large variability in size per sample is due to variability in the granularity of reference blocks. In particular, integer precision GQs produce very large files. GVCF sizes should not be compared directly to other representations to assess compression, because in many cases some fields are dropped during or after combining GVCFs.

| Description | Data type | Year | Samples | Size (TiB) |  |  | Size per sample (MiB) |  |  |
| --- | --- | --- | --- | --- | --- | --- | --- | --- | --- |
|  |  |  |  | PVCF | GVCFS | VDS | PVCF | GVCFS | VDS |
| gnomAD v2 | genomes | 2016 | 15,496 | 16.00 | 5.00 |  | 1,082.68 | 338.34 |  |
| gnomAD v3.1 | genomes | 2019 | 153,030 |  |  | 37.66 |  |  | 258.05 |
| CCDG exomes | exomes | 2021 | 203,664 |  |  | 3.36 |  |  | 17.30 |
| CCDG genomes | genomes | 2021 | 136,959 |  |  | 108.73 |  |  | 832.45 |
| gnomAD v4 | exomes | 2021 | 955,359 |  |  | 18.30 |  |  | 20.09 |
| BGE Wave 1 | exomes | 2023 | 82,000 |  | 8.76 | 2.07 |  | 112.02 | 26.47 |
| All of Us April 2023 Data Freeze | genomes | 2023 | 245,350 |  | 292.61 | 20.95 |  | 1,250.56 | 89.54 |

### Cost of joint calling

As of August 2023, using Hail 0.2.120, a VDS produced from GVCFs costs approximately 0.005 USD per exome, and in a large-scale application of the VDS combiner for whole genomes in 2019, we observed a cost of 0.10 USD per genome. These numbers represent the sum total cost of using a Google Dataproc cluster with n1-standard spot instances to generate a VDS.

### Cost of storage

The VDS is stored as a hierarchy of files or blobs. As mentioned before, the size of a VDS depends significantly on the granularity of reference blocks. As a concrete example, at 260 MiB per genome or 26 MiB per exome, a VDS costs about 0.0052 USD per genome per month or 0.00052 USD per exome per month to store in a cloud object store (at a typical cost of 0.02 USD per GiB per month).

## Conclusion

### A concise, portable, & analyzable representation

We have presented the Scalable Variant Call Representation, a concise representation of genome sequencing datasets that enables quality control and analysis of nearly one million whole exome sequences. We reified SVCR in the VCF format as SVCR-VCF. We also presented the Hail VDS, a format generated and consumed by the open-source Hail Query library.

### Joint calling

“Joint calling” has historically referred to combining samples into one dataset and also possibly adjusting genotypes given information from other samples. In this paper, we use the term “joint calling” to mirror conventional descriptions of these processes; however, “combining” may be more precise. The Hail VDS Combiner does not adjust genotypes nor is the SVCR format affected by adjustment, although notably, GATK-based “joint calling” pipelines such as GenotypeGVCFs do not do so either. Instead, SVCR and the VDS Combiner provide a representation and platform on which genotype adjustment could be implemented, but the primary challenge at scale is the transposition of single-sample GVCF files into a variant-major layout.

### Standardization

Local allele indexing has been under discussion for a few years at the hts-specs repository under issue 434 (Farjoun 2019). We hope the release of this paper motivates finalizing those changes. Further work remains to standardize reference blocks.

### Future directions

Sparse representations of genotype matrices offer an efficient way to store large-scale sequenced cohorts, but many analyses still expect dense matrices as inputs.

Developing new methods for querying and modeling genotype matrices that operate directly on sparse and/or compressed data will lead to substantial savings of compute time in addition to storage. Further, representing missing genotypes in VDS or a split SVCR format using a third table (in addition to reference and variant data) can support efficient analysis when reference quality information is not necessary by obviating the need to read the reference data; however, determining the best approach to do so in the context of dynamic filtering makes this challenging.

### Related work

*GLNexus* is both a tool and a particular modification of the PVCF format. (Lin, et al. 2018). GLNexus combines GVCFs into PVCFs in three steps: (1) find a unified set of loci, (2) “project” genotype-level fields from each GVCF into the PVCF, and (3) adjust genotype calls.

GLNexus proposes a new allele unification algorithm which introduces a new PVCF variant type: <MONOALLELIC>. This algorithm and new variant are designed to generate variants that are “completely non-overlapping, with mutually-exclusive alleles”.

GLNexus stores genotype records in a key-value store. GLNexus splits the reference genome into a disjoint set of 30 kilobase “bins”. Each genotype record is associated with a key pair of the bin containing this record and the sample identifier. GLNexus uses an LSM-tree based key-value store which permits the addition of new samples in amortized linear time and space. Genomic intervals may also be retrieved in amortized linear time.

(Lin, et al. 2019) and (Yun, et al. 2021) report the runtime and PVCF size for several applications of GLNexus to produce a PVCF from GVCFs, including an exome cohort of nearly 250,000 samples and a genome cohort of nearly 23,000 samples. We summarize their results in Table 4.

**Table 4.** GLNexus performance. We collect here the time to generate and size of PVCFs generated by GLNexus.

| Description | Reference | Samples | Time<br>(core-hours / sample) | Size<br>(MB / sample) |
| --- | --- | --- | --- | --- |
| Chromosome 2 of an exome,<br>unspecified version | Lin, et. al 2019 | 50,000 | 0.005 | Not reported. |
| Exome,<br>standalone GLNexus | Lin, et. al 2019 | 16,521 | 0.058 | Not reported. |
| Exome,<br>distributed GLNexus | Lin, et. al 2019 | 243,953 | 0.23 ("thread"-hours) | 28 |
| Genome,<br>distributed GLNexus | Lin, et. al 2019 | 22,609 | 7 | Not reported. |
| Genome,<br>unspecified version | Yun, et. al 2021 | 2,504 | Not reported. | 387 |

The innovations of GLNexus and SVCR address different PVCF challenges. SVCR, as defined above, requires one row per locus; however we believe GLNexus PVCFs are otherwise compatible with local allele indexing and preservation of reference blocks. The GLNexus database preserves reference information and effectively uses local allele indexing because it exactly preserves the GVCF records. The 30 kilobase reference bins resolve the challenge of point-queries for a dense vector in a manner similar to our “split at fixed periods” approach. The Hail VDS Combiner does not adjust genotype calls; however we believe the GLNexus empirical frequency prior could be implemented in Hail Query.

*GATK’s GenotypeGVCFs* command produces a PVCF from either one or more GVCFs or a GenomicsDB workspace. The largest callset known to the authors to use GenotypeGVCFs directly on GVCFs is the 91,796 sample ExAC unfiltered callset (Lek et al. 2016). The largest callset to use GenomicsDB is the 164,332 individuals with exome sequence data and 20,314 individuals with genome sequence data from the unfiltered gnomAD v2 callset (Karczewski et al. 2020). GenotypeGVCFs suffers from both of the problems described above: for every variant site, every sample must have a genotype and for every allele, every genotype must have quality metrics.

*GenomicsDB* (Datta 2017) and *TileDB* (Papadapoulos 2016) are projects with a shared origin which use similar techniques of sparse matrix representations to support better scaling. TileDB is an open-source multidimensional array storage manager written in C++ with support for large dense and sparse arrays. TileDB-VCF is an open-source C++ library with Python and command line interfaces which includes schemas and functionality for variant stores, including the ability to ingest new samples and query slices. TileDB-VCF stores GVCF variant and reference block records directly without PL expansion in a sparse array, indicating that

TileDB-VCF disk footprints will scale linearly in the number of samples. GenomicsDB is a separately-maintained open-source project originally developed on TileDB with the same data model described above, and tighter integration with GATK.

*The multi-sample VCF (msVCF) and DRAGEN gVCF genotyper* were developed by Illumina for generating and representing a jointly called dataset (Illumina 2023) (Schulz-Trieglaff 2023). The DRAGEN gVCF genotyper accepts either GVCFs or “multi-sample GVCFs” as input and produces an msVCF as output. This process involves three steps:

1. DRAGEN aggregates batches of GVCFs into *census files* and *cohort files*. A census file stores variant metadata and per-sample reference blocks for a batch. A cohort file is a GVCF-like format containing all the samples of a batch.
2. DRAGEN aggregates all census files into a single global census file.
3. DRAGEN uses the global census file to generate, per cohort file, an msVCF.

The first step is parallelizable per batch of GVCFs. If desired, the second and third steps can be done in a tree-like fashion which allows both scaling of samples as well as incremental addition of samples. The DRAGEN gVCF genotyper is available both on-sequencer and in the cloud.

The msVCF uses a slight variant of local allele indexing. Instead of “LA” they use “LAA” and instead of requiring and including an entry for the reference allele, they always elide the reference allele. In contrast, this paper requires the first entry of *LA* to always be 0. The msVCF defines the LPL and LGT fields equivalently to this paper. Similarly to VDS, but unlike SVCR-VCF, DRAGEN stores the reference blocks in a separate file, the census file. We believe a split version of SVCF-VCF could serve the needs of DRAGEN while also being interoperable with other tools.

DRAGEN 4.0 has generated a 500,000 sample cohort (Schulz-Trieglaff 2023). Illumina’s cloud-based DRAGEN platform, Illumina Connected Analytics, has been tested with cohorts of at least 100,000 samples. Illumina reports a cost of “0.3 iCredits per sample”. The cost of an iCredit depends on the volume of iCredits purchased, but is roughly 1 USD as of 2023-11-20. DRAGEN also supports a “compact GVCF” representation which omits statistics only necessary for *de novo* variant calling in pedigrees. This innovation is compatible with the SVCR and VDS Combiner.

*The Genomic Variant Store*, developed by the Broad Institute’s Data Sciences Platform (DSP), is a Google BigQuery based solution to GVCF aggregation and variant storage. The GVS representation is a variant of SVCR. It consists of three tables: *vet* (variants), *ref_ranges* (reference blocks), and *alt_allele* (variants). The vet and ref_ranges table are indexed by genomic position. The alt_allele table is indexed by sample. Much like GLNexus, it preserves GVCF records and thus effectively uses local allele indexing. GVS stores reference and variant data in distinct tables; indeed, it inspired the Hail VDS split representation. Storing the variant information twice, once row-major and once column-major appears unique to GVS and allows for rapid access to a single sample’s sequence. GVS can export to both PVCF and Hail VDS. As of October 2022, GVS can produce a 10,000 sample joint callset in less than half a day at a cost of 0.06 USD per genome (Degatano 2022). GVS, using Hail Query and BigQuery together, produced the All of Us April 2023 freeze callset, a 245,000 whole genome VDS.

*Other projects.* Multiple projects propose better encodings and compression of PVCF data. Savvy (LeFaive 2021) is a storage layer and C++ query API for efficient storage and queries of variant data. Savvy’s data model has the same scaling characteristics as PVCF. Lossless textual encodings of PVCF files have also been proposed. Lin et al. proposed spVCF which achieves O(N^1.1^) scaling of bytes in samples (Lin 2020). Eggertsson et al. propose popvcf which demonstrates further improvements over spVCF but they do not report a scaling factor (Eggertsson 2022). Unlike Savvy, spVCF, and popvcf, the size in bytes of SVCR and VDS both scale as O(N) in the samples. Moreover, the VDS is integrated with the scalable Hail Query dataframe and linear algebra system.

BCFTools supports local allele indexing, albeit using the *LAA* name and omitting the entry mapping the local reference allele to the global reference allele (Li 2023). This definition of *LAA* matches that of msVCF, described above.

## Acknowledgements

We thank Mike Wilson for help with HGDP data, as well as Katherine Chao, Grace Tiao, and the gnomAD consortium for making data available for methods development. We thank the rest of the Hail team and its alumni for their development of the Hail system. We thank the Data Sciences Platform (formerly Data Science and Data Engineering) broadly and especially Laura Gauthier, Yossi Farjoun, Eric Banks, and Louis Bergelson for discussion and collaboration around early prototypes. We also thank the Variants team at the Data Sciences Platform at the Broad for a productive and interesting collaboration as well as feedback on this document. We thank Lee Lichtenstein for data on the AoU VDS. This work is supported by 1U01MH115727-01, 1UM1HG008900-01, U24HG011450-03, 1U01MH115727-01, 5U54DK105566-03, 2R01MH107649-04, 5U01HG009088-03, U24HG011450, the Chan Zuckerberg Initiative, as well as The Stanley Center for Psychiatric Research, which together with the Neale Lab has provided an incredibly supportive and stimulating home. This work is also supported by the Novo Nordisk Foundation (NNF21SA0072102) and R37MH107649. We thank the Jeremy M. and Joyce E. Wertheimer Foundation, whose strategic advice and generous philanthropy have been essential for growing the impact of Hail. The content is solely the responsibility of the authors and does not necessarily represent the official views of any funding source.

